# Soil microbial composition varies in response to coffee agroecosystem management

**DOI:** 10.1101/2020.07.10.197202

**Authors:** Stephanie D. Jurburg, Katherine L. Shek, Krista McGuire

## Abstract

Soil microbes are essential to the continued productivity of sustainably-managed agroecosystems. In shade coffee plantations, the relationship between soil microbial composition, soil nutrient availability, and coffee productivity have been demonstrated, but the effect of management on the composition of the soil microbial communities remains relatively unexplored. To further understand how management modulates the soil microbiome, we surveyed the soil fungal and bacterial communities, as well as soil chemistry and canopy composition in a Nicaraguan coffee cooperative, across 19 individual farms. Using amplicon sequencing, we found that management (organic or conventional), stand age, and previous land use strongly affected the soil microbiome, albeit in different ways. Bacterial communities were most strongly associated with soil chemistry, while fungal communities were more strongly associated with the composition of the canopy and historical land use of the coffee plantation. Notably, both fungal and bacterial richness decreased with stand age. In addition to revealing the first in-depth characterization of the soil microbiome in coffee plantations in Nicaragua, our results highlight how fungal and bacterial communities are simultaneously modulated by long-term land use legacies (i.e., an agricultural plot’s previous land use) and short-term press disturbance (i.e., farm age).

**One-sentence-summary:** The composition of soil fungal and bacteria communities in shade coffee plantations depend on the combination of the farm’s management type, its previous land use, and the coffee plants’ stand age, but are differently influenced by each.

## INTRODUCTION

The biotic and abiotic drivers of microbial community assembly across space and time determine the composition of a local community, which in turn determines the functional properties of that community (Nemergut *et al*. 2013). These linkages are especially critical to examine in human-modified terrestrial ecosystems, as microbes perform critical functions related to the resistance and resilience of aboveground plant communities to stress and disturbance (Wardle *et al*. 2004; van der Putten *et al*. 2016), and drive the vast majority of biogeochemical cycling in soils (Six *et al*. 2006; Falkowski, Fenchel and Delong 2008). Agriculture is one of the most widespread and significant ecosystem modifications by humans, and cultivated land now accounts for 36.6% of the terrestrial surface (FAO 2017). Consequently, determining the assembly mechanisms of microbial communities in agroecosystems is essential for leveraging their benefit for crop production, pest management, and soil function (Brussaard, De Ruiter and Brown 2007; Berg 2009; Chaparro *et al*. 2012; Toju *et al*. 2018).

Coffee plantations are important agroecosystems for evaluating the role of microbial assembly mechanisms, as they are one of the most highly traded commodities in the world (Donald 2004; Ricketts *et al*. 2004) and have significant variation in the intensity by which they are managed (Philpott *et al*. 2008; Tscharntke *et al*. 2011; Perfecto, Vandermeer and Philpott 2014). In Central America, coffee production constitutes an important part of local economies, and coincides with areas of extremely rich biodiversity (Komar 2006). Coffee is especially important for the Nicaraguan economy, where exports of the crop make up approximately one fifth of the country’s exports (Bacon *et al*. 2008). Following the proliferation of industrial coffee plantations and the extensive clearing of forests for agriculture in Nicaragua in the 1960’s-1980’s, agrarian reform stimulated the growth of Fair Trade cooperatives (Bacon 2010). International concern about the effects of deforestation on the biodiversity of the region coupled Fair Trade efforts with conservation initiatives, stimulating the creation of shade coffee plantations as faunal reserves (Komar 2006), as well as for the maintenance of ecosystem services (Goodall, Bacon and Mendez 2014; Perfecto, Vandermeer and Philpott 2014).

In shade coffee plantations, coffee (*Coffea arabica L*.) is cultivated, and tree cover is managed to provide shade for the crops. Canopy tree composition may be an important biotic filter for soil microbial communities due to known aboveground-belowground linkages in many terrestrial ecosystems including tropical forests (Peay, Baraloto and Fine 2013; Barberán *et al*. 2015). Cover crop composition is also known to affect microbial composition and function in other agricultural systems (Vukicevich *et al*. 2018; Hallama *et al*. 2019; Ma *et al*. 2020), so it is likely that this also occurs in coffee plantations. Shade cover has been associated to increased coffee market quality (Vaast *et al*. 2006), as well as higher organic matter content and reduced nutrient leaching (Babbar and Zak 1995; Beer *et al*. 1997). In practice, canopies in coffee agroecosystems are managed in a highly individualized manner (Hernández-Martínez, Manson and Hernández 2009). The canopy overlying shade coffee plantations may be managed to provide the farmer with additional resources such as timber and fruits. The resulting compositional patchwork likely affects the soil microbiota to different extents. Shade trees in agroecosystems also provide other important ecosystem services, including pollination, pest control, and increased connectivity between forest patches in the vicinity of agroecosystems (Philpott *et al*. 2008; Goodall, Bacon and Mendez 2014; Perfecto, Vandermeer and Philpott 2014). The effect of shade coffee management has assessed for ants (Philpott 2005), birds (Buechley *et al*. 2015), epiphytes (Goodall, Bacon and Mendez 2014), and woody plants (Kumsa *et al*. 2016), but few studies have explicitly studied soil microbial communities. To date, studies of soil biota under shade coffee have found that soil carbon associated with microbial biomass is consistently higher in organically-managed than in conventional, intensively-managed coffee plantations (Partelli *et al*. 2012), and is associated with higher coffee yields (Silva Aragão *et al*. 2020). Shade trees can also increase soil N and the abundance of ammonia oxidizing bacteria, which convert ammonia into nitrite in the process of nitrification (Munroe *et al*. 2015). Together, these studies suggest that management practices, including shade tree management, may directly influence the assembly of soil microbial communities in coffee agroecosystems.

Here, we explored the relationship between management practices, canopy tree composition, soil biochemical composition, and soil fungal and bacterial communities within Nicaraguan agricultural cooperatives. Our study targeted 19 coffee farms in Sontule, a small cooperative in the Northern Pacific Nicaraguan highlands. The farms varied in time from conversion to coffee plantations, canopy management strategies, and coffee plant age, which may affect soil microbial communities due to legacy effects (Kulmatiski and Beard 2008; Fichtner *et al*. 2014; Schrama *et al*. 2016). By providing an in-depth examination of how management strategies and the resulting tree composition and soil chemistry modulate the soil microbiome, our study informs future shade management strategies.

## MATERIALS AND METHODS

### Study site and sample collection

The Unión de Cooperativas Agropecuarias (UCA) Miraflor, is a union of small agricultural cooperatives located within the Miraflor Nature Reserve, 30 km from the city of Estelí, Nicaragua (13.25966,-86.243711, Fig. S1, Supporting Information). The UCA consists of approximately 70 farmers who manage several plots of coffee with varying degrees of shade and management intensity. Management strategies were divided into conventional management, which included inputs of herbicide and industrial fertilizer; and organic management, in which soil inputs were limited to various composts. In July 2011, 19 coffee plots were selected after interviewing farmers about their management strategies, age of the plantation, previous land use of the plot, and age of the current coffee plants in order to obtain a representative subset of the UCA (Table S1, Supporting Information). These included plots with a range of coffee plant ages (1:45 years, mean= 12 years) and previous land uses (primary forest, n=5; secondary forest, n=7; coffee plantations, n=6). Five of the plots were conventionally managed, with little or no shade coverage. In addition to the 19 plots, two unmanaged areas with primary and secondary forest were sampled. All plots were at an elevation of between 1018 and 1267 m above sea level (mean 1182 m).

For each plot, three 5×5 m replicate subplots were randomly selected. For each subplot, canopy cover was assessed with a densiometer, measured 4 times per subplot and averaged. All trees whose canopy fell within the subplot and with diameter breast height greater than 10 cm were identified by their local name and listed. From each subplot, triplicate 5 cm cores were randomly sampled from the top 10 cm soil layer. Triplicate cores from each subplot were later homogenized by sieving through a 2 mm sieve. Soil element analyses were performed at the Auburn University Soil Testing Laboratory (AL, USA), using inductively coupled plasma atomic emission spectroscopy. Soil C and N were measured using loss on ignition.

### DNA extraction, amplification and sequencing

DNA was extracted using the MoBio PowerSoil extraction kit (MoBio, Carlsbad, CA), according to the manufacturer’s instructions, with modifications as previously described (Hobbs *et al*. 2006; Lauber *et al*. 2009). Bacterial samples were sequenced as part of the Earth Microbiome Project (EMP, Thompson et al., 2017), and fungi were sequenced at the New York Medical College Genomics Core Laboratory (Valhalla, NY). Bacterial communities were assessed by amplifying and sequencing the V3-V4 region of the 16S rRNA gene with primers 515F and 806R (Caporaso *et al*. 2012), and fungal communities were assessed by amplifying the internal transcribed spacer (ITS1) region with primers ITS1-F and ITS2 (McGuire *et al*. 2013). Both sets of amplicons were sequenced on an Illumina Hiseq with a paired-end 150-bp sequencing kit. Raw sequences were demultiplexed and processed using QIIME and the UPARSE pipeline (Edgar 2013). Sequences were clustered into operational taxonomic units (OTUs) at a 97% similarity threshold and rarefied to 7000 and 20700 sequences per sample for bacteria and fungi, respectively. Sequences were then assigned taxonomy using BLAST, prior to downstream analysis. Only samples for which both fungal bacterial sequences were available were preserved for analyses, resulting in a final dataset which described 19 farms with 48 samples in total (Supporting Information). Bacterial data are publicly available as part of the Earth Microbiome Project (Thompson *et al*. 2017), and fungal data is available in the Supporting Information, and will be deposited into a public repository upon manuscript acceptance.

### Statistical analyses

We evaluated α-diversity as the number of fungal and bacterial OTUs per sample (species richness). Tree richness was measured as the number of different species whose canopy fell within a subplot. Correlations were evaluated with Pearson’s R. Values are presented as *mean* ± *sd* throughout the manuscript. Differences in means were tested non-parametrically with Wilcoxon rank sum tests for comparisons between two groups, and with Kruskal-Wallis tests for comparisons between more groups.

Multivariate analyses were performed with the vegan (v.2.5-6) and phyloseq (v.1.26.0) packages (Oksanen *et al*. 2007; McMurdie and Holmes 2013). Fungal, bacterial, and canopy tree composition as well as soil chemical composition were Hellinger-transformed and converted to Bray-Curtis distances prior to all analyses. The significance of all multivariate analyses was evaluated by permutation tests (999 iterations).

The relationship between fungal, bacterial, and canopy tree composition, as well as soil chemistry was evaluated in pairwise Mantel tests. The effect of management parameters (Organic/conventional management, the farm’s previous land use, and the age of the coffee plants) on fungal and bacterial community composition was evaluated with PERMANOVA-like adonis tests. To investigate the effect of management practices on soil chemistry, we performed a distance-based RDA (db-RDA) on the soil chemistry matrix, constrained by the three management parameters. Interactive terms were not included. The significance of the model, the explanatory terms, and each axis were evaluated by permutations (*anova*.*cca*). To further break down the relative contribution of each management parameter, we performed a variance partitioning analysis with the same terms (*varpart*). To evaluate the effect of management, canopy composition, and soil chemistry on the soil fungal and bacterial communities, we performed backwards and forwards selection of explanatory variables (*ordiR2Step*, perm.max=200). For each community, variables which were selected were included in the db-RDA model, evaluated as previously.

Putative functional guilds were assigned to fungal OTUs using the FUNGuild program (Nguyen *et al*. 2016). To evaluate the effects of management on functional guilds of soil fungi, OTUs assigned as plant pathogens and endophytes were subsetted and agglomerated at the genus level to be analysed separately. Shifts in relative abundances of fungal functional groups across management and age of coffee plants were evaluated using PERMANOVA. We used principal component analysis (PCA) to construct axes of edaphic variation using the following soil chemical variables: Ca (ppm), K (ppm), Mg (ppm), P (ppm), Al (ppm), Ba (ppm), Fe (ppm), Mn (ppm), Na (ppm), Zn (ppm), B (ppm), Cu (ppm), total %C, total %N, and moisture content (%). Relationships between abiotic soil parameters and fungal plant pathogen and endophyte communities were visualized using the first principal component axis and relative abundance data. All data visualization and figures were generated using ‘ggplot2’ (Wickham 2011).

## RESULTS

Average canopy cover did not differ between organically and conventionally-managed farms, however, organically-managed farms had a significantly higher canopy tree richness than conventionally-managed farms (5.94±1.79 and 3.42±3.02 trees per plot, respectively; p=0.001). Soil moisture was highest in plots converted from primary forest, and lowest in plots converted from previous coffee plantations (χ^2^ =8.52, df=2, p=0.014).

On average across all samples, soil bacterial communities were dominated by Proteobacteria (32.4±3.7% of the community), Verrucomicrobia (17.9±5.4% of the community), and Acidobacteria (16.5±2% of the community). Organically-managed plots had consistently higher relative abundances of WS3 (Latescibacteria), while conventionally-managed plots had higher relative abundances of Nitrospirae and Chloroflexi (t-tests, p<0.05 for all comparisons, Fig. 1). Fungal communities were dominated by Ascomycota (37.2±15.5% of the community), Zygomycota (34.6±20.2% of the community), and Basidiomycota (11.9±11.2% of the community). A large portion of the fungal community (12.3±6.7%) was unclassified at the phylum level, and organically-managed plots had a higher relative abundance of Ascomycota, while conventionally-managed plots had higher relative abundances of Chytridiomycota (Fig. 1). Fungal communities were more variable across plots than bacterial communities (Fig. S2, Supporting Information).

**Figure 1.**
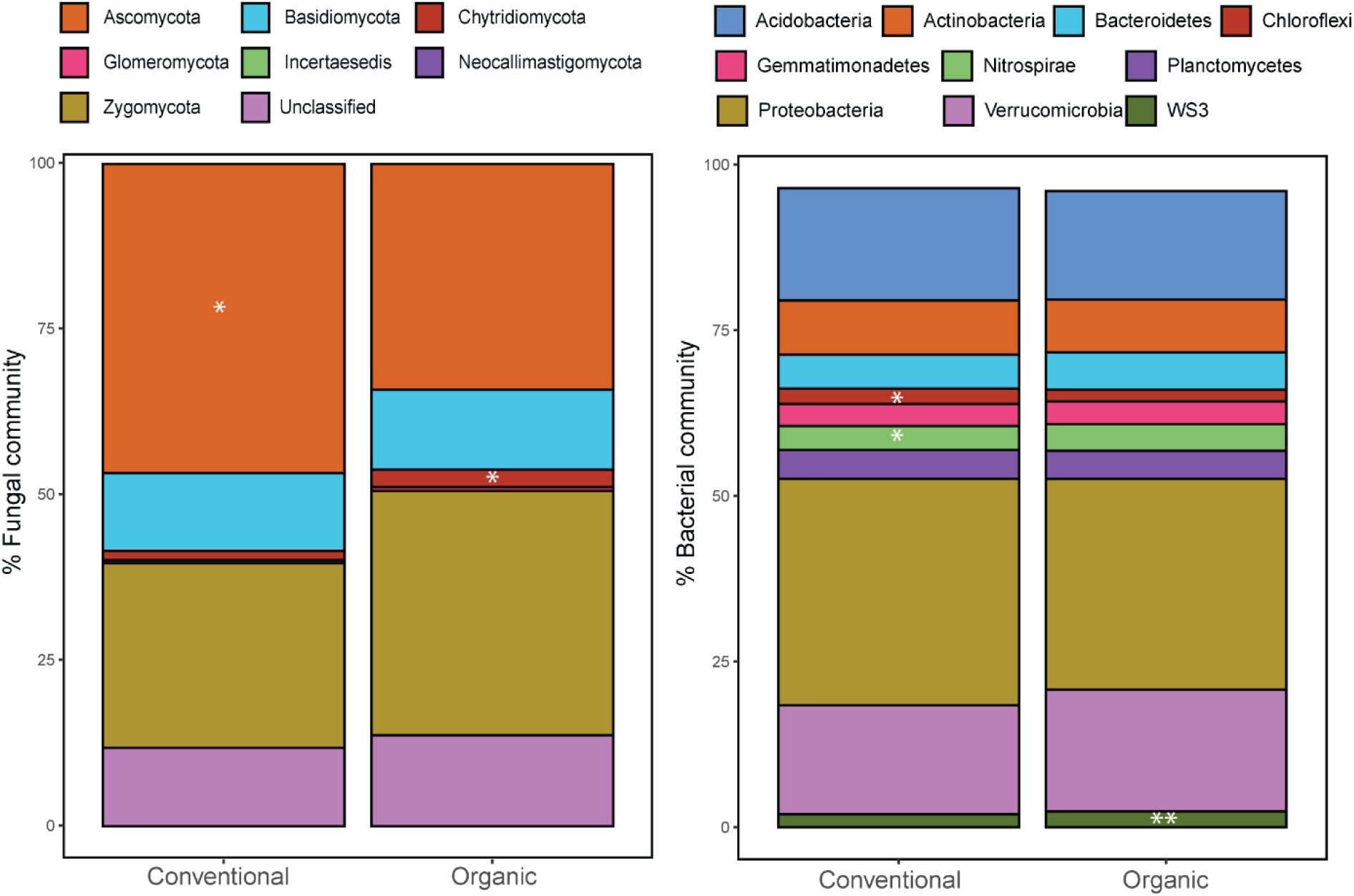
Relative abundance of the dominant fungal (left) and bacterial (right) phyla on average, for each management type. All fungal phyla are represented. For bacteria, the 10 most dominant phyla on average across all samples are displayed. White asterisks indicate significantly higher relative abundances in a phylum for that management type, as determined by t-tests (*** p<0.001; **p<0.01; * p<0.05). The relative abundances of individual samples are available in S2.

To examine how farm-specific management practices affected soil chemistry, we performed a db-RDA on the soil chemical composition, with organic/conventional management, previous land use, and the age of the coffee plants as explanatory variables This model explained 27.4% of the variance in soil chemistry and was significant (*p*=0.001), as were the three terms added (p≤0.01, Fig. 2). Whether the farm was managed organically or conventionally was the largest driver of soil chemistry (8.2%), followed by the farm’s previous land use (7.5%) and the age of the farm’s coffee plants (3%).

**Figure 2.**
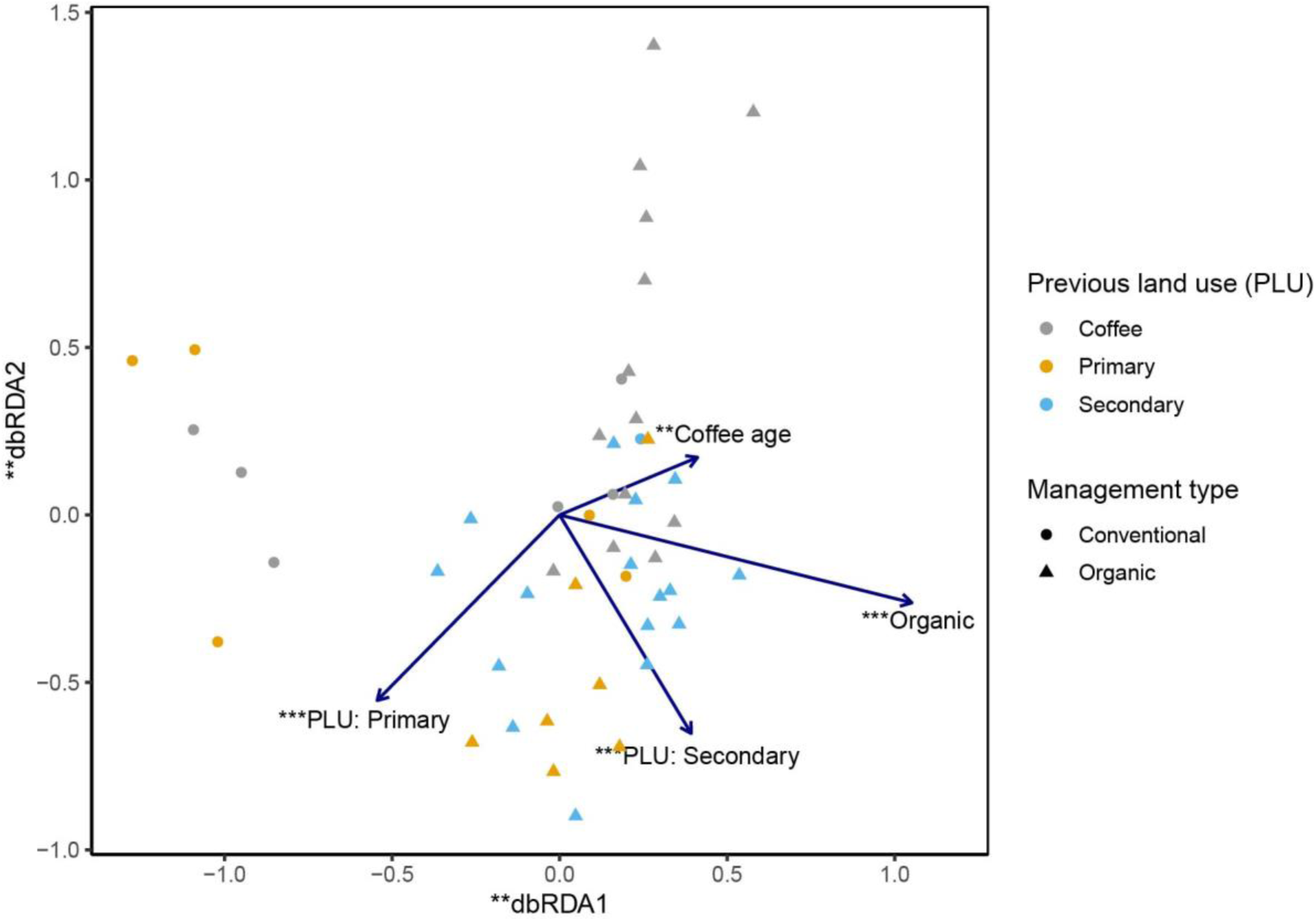
A db-RDA shows how different aspects of management affect soil chemistry. Arrows corresponding to conventional management and a previous land use are omitted as they are linearly dependent on the other levels of their respective variables. Circles and triangles represent plots. The db-RDA was significant (p<0.001), and the significance of each term and axis is indicated with asterisks (*** p<0.001; **p<0.01; * p<0.05).

Different aspects of management affected the concentration of specific elements in soil. Organically managed farms had higher percentages of carbon and nitrogen, while conventionally-managed farms had higher concentrations of phosphorus (p<0.05 for all Wilcoxon rank sum tests). In contrast, zinc, aluminium, and manganese were significantly higher in farms that were previously coffee plantations, while sodium, carbon, and nitrogen had higher concentrations in lands that were converted from secondary forest, and calcium was highest in lands that were converted from primary forests (p<0.05 for all Kruskal-Wallis tests). Coffee plant age was negatively correlated with potassium and calcium (R=-0.31 and - 0.31 respectively; p<0.05 for all correlations), and positively correlated with zinc, (R=0.33, p=0.02).

### Alpha diversity

We evaluated how management practices affected the number of bacterial and fungal taxa (richness) in each plot. On average, we detected 680.5±119.8 fungal taxa and 1931±83 bacterial taxa per plot. We found no difference in fungal or bacterial richness between conventionally and organically managed farms (*p*=0.37 and *p*=0.75, respectively). In contrast, the farm’s previous land use significantly affected fungal richness (χ^2^=7.94, df=2, *p*=0.019), but not bacterial richness (χ^2^=0.17, df=2, *p*=0.92; Fig. 3A, 3B). Farms which had been converted from primary forest had lower fungal diversity (574.7±65.3 taxa) than those which had been converted from secondary forest or long-term coffee farms (718.3±112.7 taxa). Among organically managed farms, both fungal and bacterial richness were negatively and correlated to the age of the coffee plants in the plot, but this was only significant for bacterial richness (R=-0.73, *p*=0.007, R^2^=0.53, Fig. 3D). Neither fungal nor bacterial richness in organic farms were significantly correlated with tree canopy richness, however both fungal (R=0.37, *p*=0.03, R^2^=0.14) and bacterial (R=0.37, *p*=0.03, R^2^=0.14) richness among organic farms were positively related to average canopy cover (Fig. 3E, 3F). Bacterial and fungal richness were weakly correlated to each other (R=0.28, *p*=0.053, R^2^=0.08).

**Figure 3.**
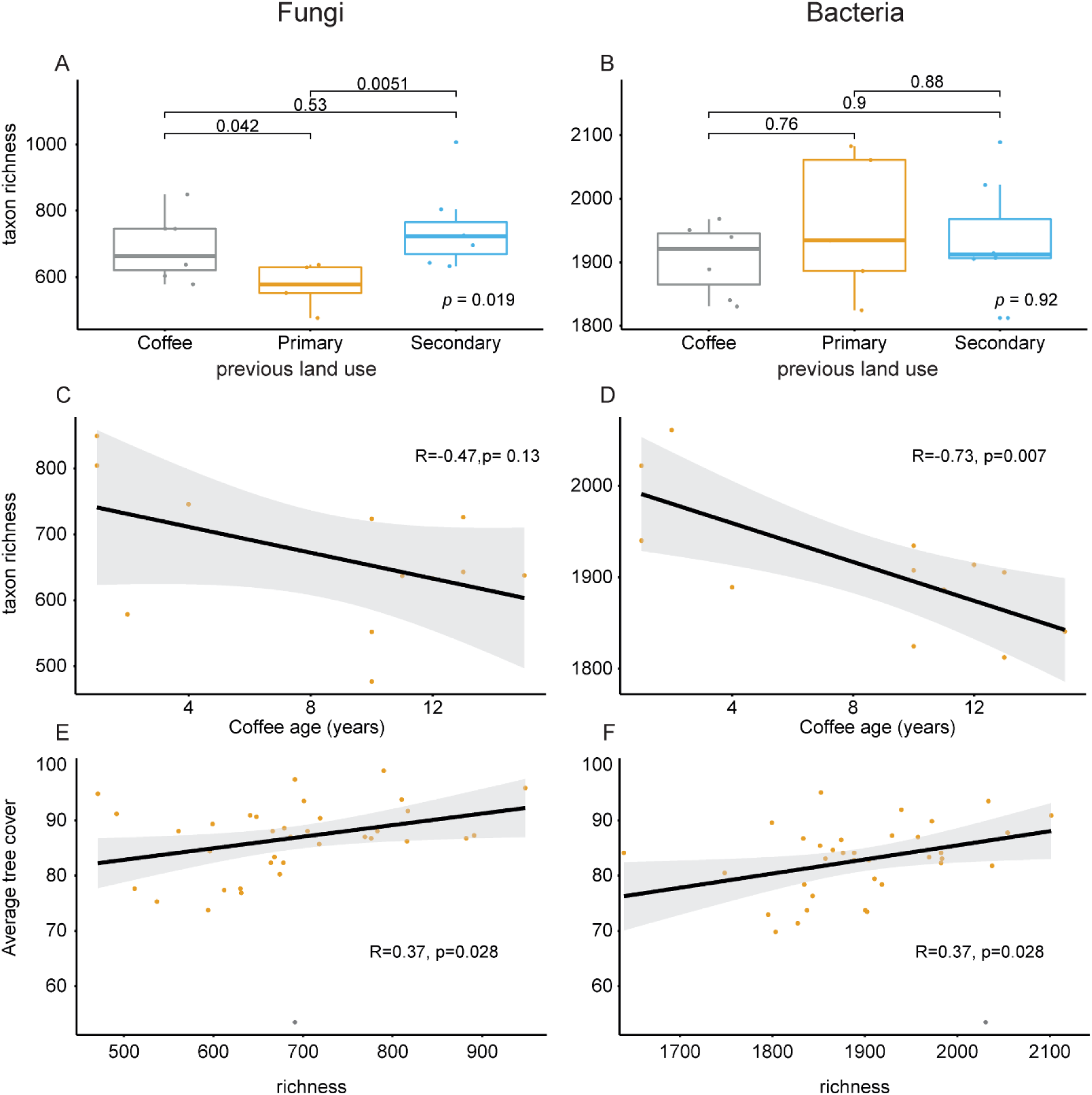
Fungal richness is significantly higher in farms converted from previous coffee farms or secondary (i.e., disturbed) forest than in farms converted from primary (i.e., old growth) forest (A), but bacterial richness is not affected by previous land use (B). In organically managed farms, both fungal (C) and bacterial (D) richness were negatively correlated to the age of the coffee plants, and positively correlated to average tree cover (E and F, respectively). Circles in A-D are the average values for the farm. Kruskal-Wallis tests in A-B were used to determine global significance (bottom left of each panel), and Wilcoxon rank sum tests were used to test pairwise comparisons. In C-F, pearson correlations were evaluated. The black line indicates a linear regression, and the grey shaded area is the 95% confidence interval for the estimate.

### Beta diversity

We assessed compositional differences in the fungal and bacterial communities across farms by looking at the Bray-Curtis distances between samples. Fungal and bacterial communities were strongly related to each other (Mantel’s r= 0.466, *p*=0.001). Adonis analysis showed that both fungal and bacterial communities clustered significantly according to whether they were organically or conventionally managed; the farm’s previous land use, and the age of the coffee plants (*p<*0.01 for all comparisons, Table 1). Among organic farms, bacterial community composition was related to the soil’s chemical composition (Mantel’s r= 0.33, *p*=0.003). In contrast, fungal communities were significantly related to tree composition (Mantel’s r= 0.17, *p*= 0.03).

**Table 1.** Results of an adonis analysis. Fungal and bacterial community matrixes were Hellinger-transformed to Bray-Curtis distances prior to analyses. The models did not include interactive terms, and significance was confirmed with 999 permutations.

|  |  | Df | SumOfSqs | R2 | F | Pr(>F) |
| --- | --- | --- | --- | --- | --- | --- |
| Fungi | Previous land use | 2 | 0.8652 | 0.05593 | 1.356 | 0.001 |
|  | Management | 1 | 0.4338 | 0.02804 | 1.3597 | 0.005 |
|  | Coffee age | 1 | 0.4512 | 0.02917 | 1.4142 | 0.001 |
|  | Residual | 43 | 13.7185 | 0.88686 |  |  |
|  | Total | 47 | 15.4686 | 1 |  |  |
| Bacteria | Previous land use | 2 | 0.3455 | 0.05619 | 1.3711 | 0.004 |
|  | Management | 1 | 0.1861 | 0.03027 | 1.4775 | 0.006 |
|  | Coffee age | 1 | 0.1996 | 0.03246 | 1.5844 | 0.005 |
|  | Residual | 43 | 5.4175 | 0.88107 |  |  |
|  | Total | 47 | 6.1488 | 1 |  |  |

To disentangle the drivers of the microbial community, we first used backwards and forwards selection to choose the most relevant tree canopy species, soil chemical components, and management parameters for fungal and bacterial communities. For fungal communities, we identified calcium, phosphorus, potassium, iron and soil carbon as relevant chemical parameters; *Rinorea squamata, Psidium spp*., *Manilkara zapota*, and an unidentified species locally known as “Cocotillo” as relevant canopy tree species; and the farm’s previous land use as a relevant management parameter. An RDA with these explanatory variables accounted for 32.4% of the variance in community composition. All explanatory variables were significant (*p*<0.05 for all terms; *p*=0.001 for the whole model, Fig. 4A).

**Figure 4.**
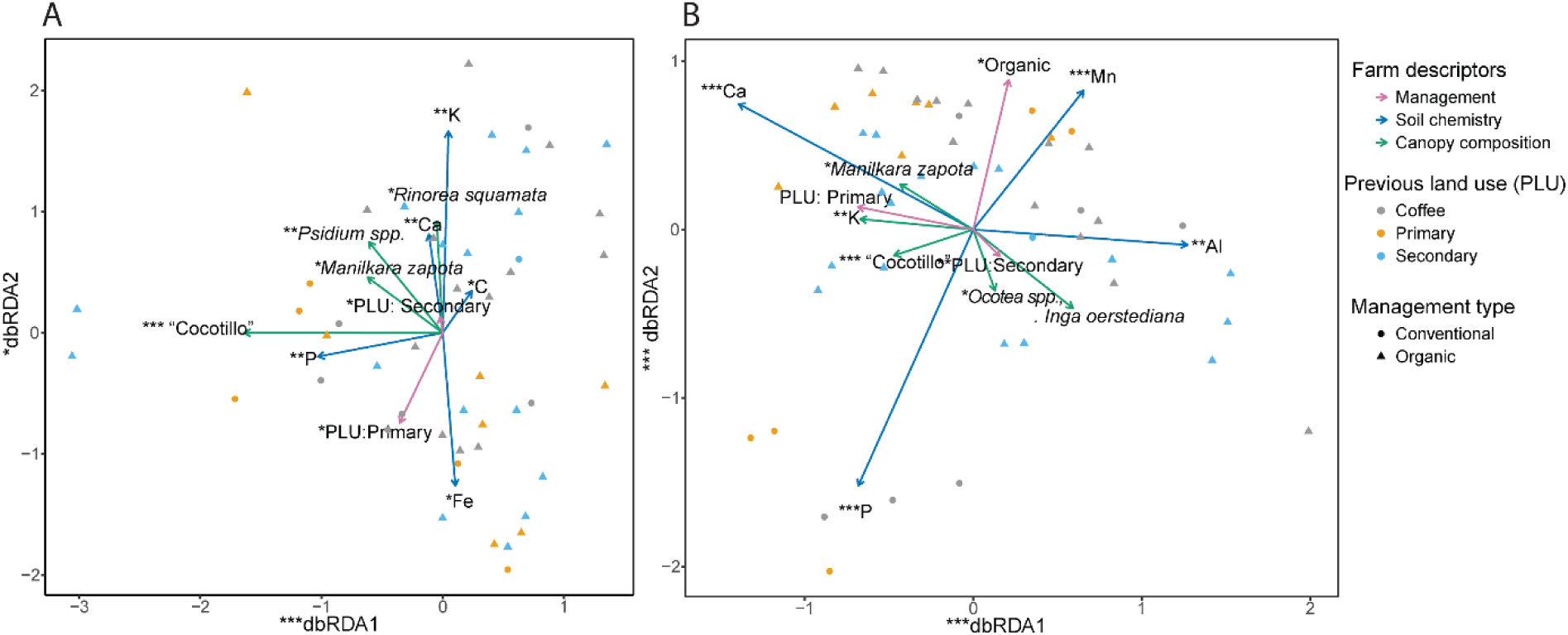
A db-RDA of selected explanatory variables shows how different aspects of management affect soil fungal communities (A) and bacterial communities (B). Arrows for explanatory variables are colored according to the type of farm descriptor; the arrows corresponding to coffee as a previous land use (both panels) and conventional management (bottom panel) are omitted as they are linearly dependent on the other previous land use types. Circles and triangles represent plots. The db-RDA was significant (p<0.001), and the significance of each term and axis is indicated with asterisks (*** p<0.001; **p<0.01; * p<0.05). The tree species locally known as “Cocotillo” could not be identified.

For the bacterial communities, backwards and forwards selection identified calcium, phosphorus, manganese, aluminium, and potassium as relevant chemical parameters; *Manilkara zapota, Ocotea spp*., *Inga oerstediana*, and “Cocotillo” relevant canopy tree species; and the farm’s previous land use and whether it was organically or conventionally managed as relevant management parameters. An RDA with these explanatory variables accounted for 36.9% of the variance in the bacterial community composition. All explanatory variables were significant (*p*<0.058 for all terms; *p*=0.001 for the whole model, Fig. 4B).

### Fungal functional guilds

Endophytes were the most abundant classified fungal functional group in conventional (22.9%) and organic (28.6%) coffee farms, while endophytes only comprised 13.3% of fungal communities in forest soils (Fig. 5A). Relative abundances of fungal plant pathogens varied with management as well, comprising 3.6% of the fungal community in conventional farms, and only 2.0% and 1.3% of organic and forest communities, respectively (Fig. 5A). When comparing functional guild composition across farms varying in time since established, endophytes and saprotrophs were the most abundant classified functional groups, with shifts in relative abundances with time since cultivation (Fig. 5B). To further illustrate the effects of management on fungal functional guilds, we examined plant pathogen and endophyte community composition and relative abundances. Fungal plant pathogen community composition shifted across management, but with minor significance (PERMANOVA p=0.085, R2=0.052, F model = 1.413); whereas, endophyte community composition did not shift significantly across management (PERMANOVA p=0.271, R2=0.043, F model=1.116) (Fig. 6A and B, respectively). Both fungal endophytes and plant pathogen composition shifted significantly across sites varying in previous land use (endophyte: PERMANOVA p=0.022, R2=0.063, F model=1.747; pathogen PERMANOVA p=0.001, R2=0.082, F model=2.337). Endophyte communities were dominated by *Pleurotus* and *Rhizopycnis* species across all samples regardless of management or previous land use.

**Figure 5.**
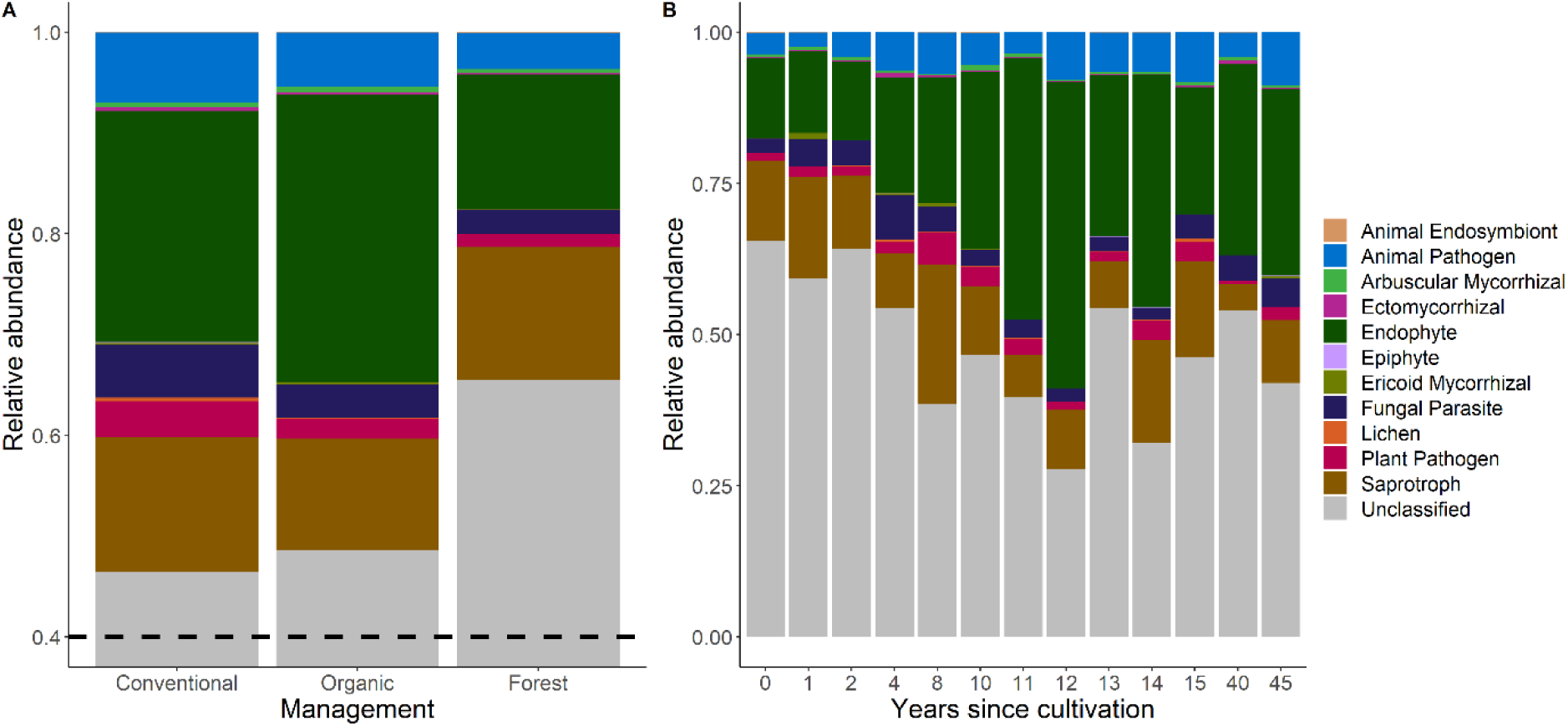
Relative abundances of functional guilds of fungi across management (A) and years since cultivation (age of coffee plants, B) as assigned by the FUNGuild program. Dashed line in panel A denotes relative abundance of 0.4, shifting the y-axis because >40% of fungal sequences across management categories were unclassified by FUNGuild.

**Figure 6.**
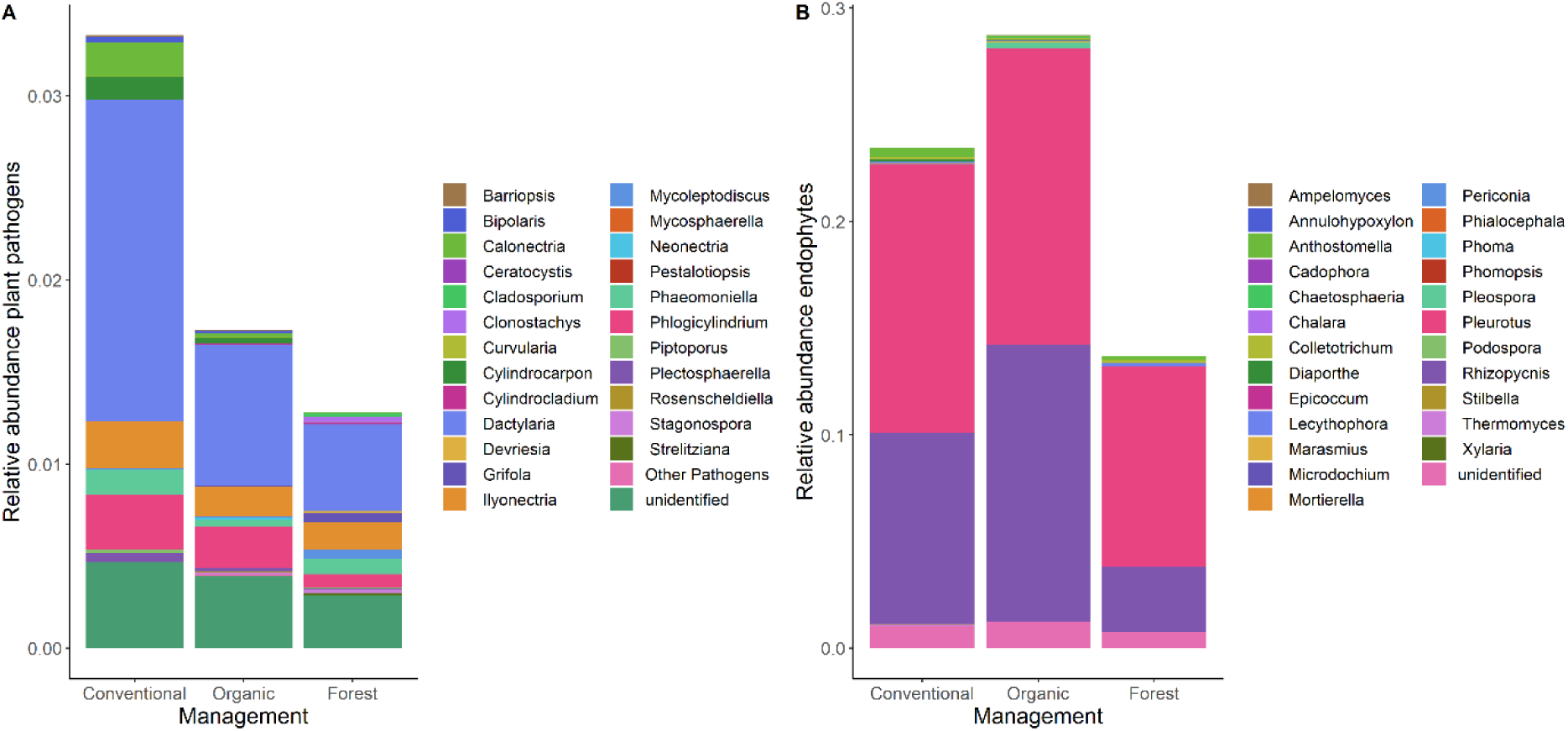
Abundances of fungal genera assigned as plant pathogens (A) and endophytes (B) relative to the whole fungal communities detected in samples divided by management category.

Principal component analysis condensed soil parameters into axes representing gradients of macronutrient content, soil C and N, and soil moisture; PC1 explained 31.11% of variation in soil chemistry across samples and runs from high mineral content (namely Fe, Cu, B, P, Al and Zn; negative values) to higher moisture and %C and N (positive values, Fig. S3). There was a significant effect of the first principal component on the relative abundances of plant pathogens (R2 = 0.19, p< 0.001), with a decline in pathogen abundance along the first PC axis (Fig. 7).

**Figure 7.**
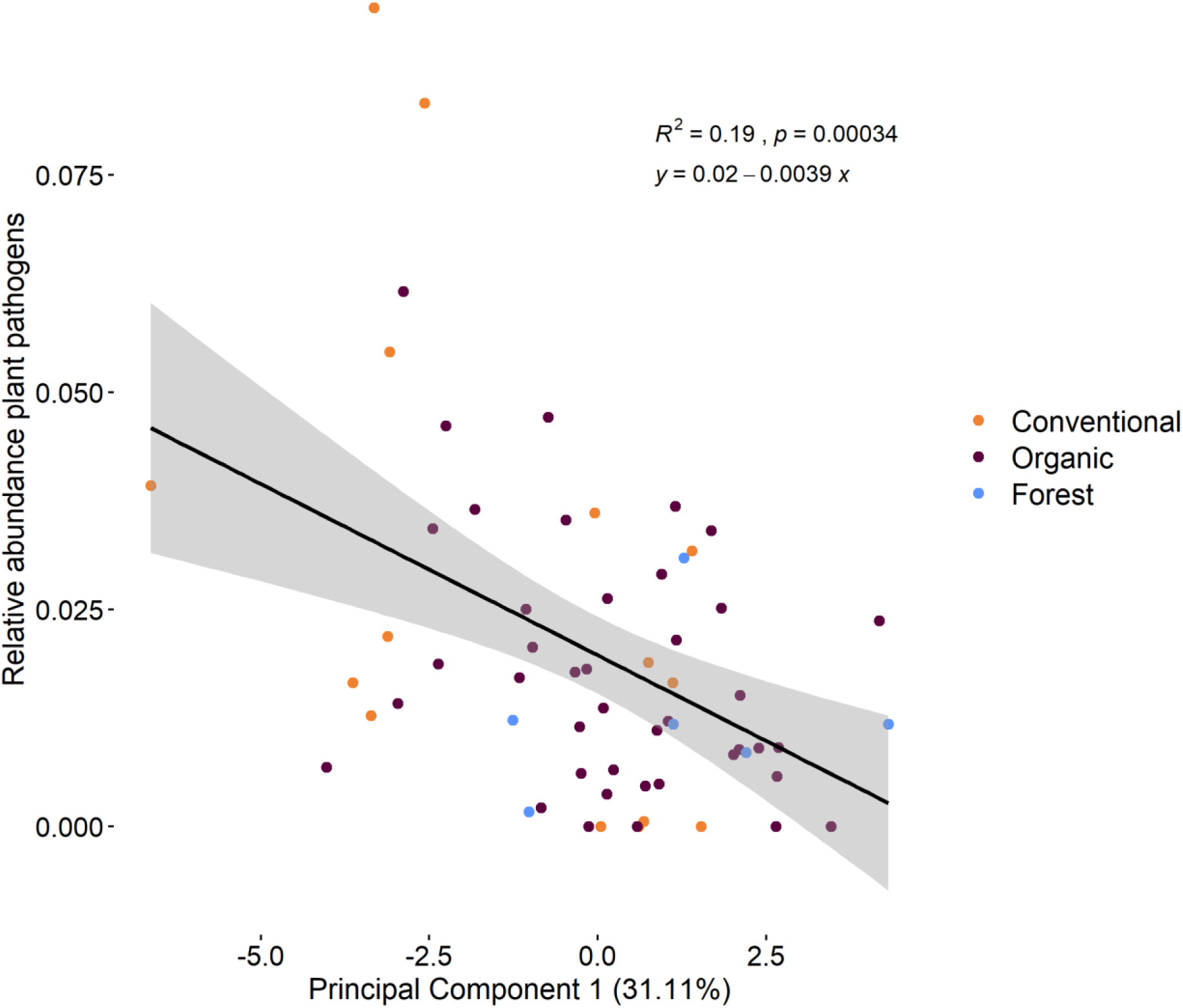
Significant regression of relative abundances of plant pathogens against the first principal component axis constructed from soil chemical parameters. Points represent individual samples highlighted by management. PC1 runs from high mineral content (Fe, Cu, B, P, Al and Zn) to higher soil moisture, %C and %N. Principal component biplot included in supplementary information.

## DISCUSSION

By analysing soils from 19 Nicaraguan coffee plantations subjected to a range of management regimes, we found that soil bacterial and fungal communities were altered by various aspects of management, which may have cascading effects on productivity, Previous studies have also found significant effects of coffee management intensity on microbial composition (Hartmann *et al*. 2015). However, we uncovered an added layer of complexity to microbial assembly processes across management intensity gradients, which was previous land use of a plantation. The historical land use of a plantation affected the fungal and bacterial compartments in different ways, but demonstrated that agricultural intensification can interact with land use legacies in a given site to modulate soil microbial communities. The age of the coffee plantation was also a significant factor explaining variation in microbial composition, suggesting that assembly processes are dynamic and have temporal variability that cannot be inferred from sampling a farm at a single time point. Furthermore, it may take longer time periods of management changes to see parallel shifts in soil microbial communities.

In conventionally-managed farms, soils received inputs of herbicides (i.e., Gramoxone) and industrial fertilizers (i.e., 15%N 15%P 15% K), while organically-managed plots received annual inputs of compost, bokashi, or agricultural pulp. Correspondingly, we found a higher abundance of carbon and nitrogen in organically managed soils, and higher phosphorus in conventionally-managed soils, in accordance with other studies of coffee agroecosystems in other countries (Partelli *et al*. 2012). Differences in soil carbon, phosphorus, and nitrogen are strong determinants of bacterial community composition (Delgado-Baquerizo *et al*. 2017) and may explain how management affected the microbiomes. Contrary to the prevailing notion that below- and aboveground richness are related (De Deyn and Van Der Putten 2005), we found no relationship between fungal or bacterial richness and canopy richness. However in our study, other farm characteristics, including previous land use and the coffee stand age, were stronger determinants of the soil microbiome.

The previous land use of each farm had a strong effect on the soil microbiome, consistent with experimental findings of legacy effects on the assembly of soil microbial communities (Jurburg *et al*. 2017; Peerawat *et al*. 2018; Turley *et al*. 2020). Our study further shows the persistent effect of previous land uses on lands under active cultivation. Fungal richness was strongly dependent on the land’s previous land use, and was lowest in farms which had been converted from primary forest, suggesting that many forest-associated taxa do not persist in converted coffee plantations. In contrast, we found no relationship between bacterial richness and previous land use. The higher sensitivity of fungal richness to agricultural land use change has been experimentally shown in restored agricultural fields compared to restored non-agricultural lands (Turley *et al*. 2020). Long-term (∼50 years) legacy effects of land use change on fungal communities have also been reported in tropical forests converted to oil palm plantations (McGuire *et al*. 2015) as well as in harvested temperate forests (Hartmann *et al*. 2015). In contrast, a meta-analysis of soil bacterial responses to land use change in the tropics found that bacterial alpha diversity increases following conversion to agricultural land (Petersen, Meyer and Bohannan 2019), possibly due to bacterial communities tracking changes in soil properties such as pH rather than responding directly to plant diversity loss following agricultural conversion.

Our results further support that fungal communities are more sensitive to land use change than bacterial communities, possibly due to the fact that fungi tend to form specialized associations with specific tree species, through differences in root exudates or leaf litter (De Deyn and Van Der Putten 2005; Prescott and Grayston 2013; Van der Putten *et al*. 2013; Barberán *et al*. 2015). Fungal communities respond more strongly than bacteria to changes in root exudates that result from altered plant diversity (Eisenhauer *et al*. 2017), which often results from land use change. Indeed, we found that fungal community composition was strongly related to tree canopy composition in coffee agroecosystems. In our study, the legacy effects of previous land use held the strongest explanatory power for soil moisture and soil chemistry, reflecting how prior land use results in long-term alterations to the soil environment, which may directly or indirectly affect microbial community composition.

The richness of bacteria, in contrast, was more strongly affected by the age of the coffee plants than other factors analysed in this study, decreasing linearly with coffee age. A similar, negative relationship between plant age and soil bacterial richness was reported for rubber trees (Peerawat *et al*. 2018), but to our knowledge, this experiment is the first to report such a relationship in the coffee agroecosystem. Slight shifts in management frequency (Peerawat *et al*. 2018) and changes in plant phenology with stand age may explain this pattern, but these two are interlinked and cannot be easily resolved, as management often depends on phenology. The production of coffee beans begins approximately three years after seed germination, and while the plant may stay productive for up to 80 years, the economic lifespan of the *C. arabica* plant is approximately 30 years (Wintgens 2009). In our study, the oldest plants were 45 years of age, after which farmers reported cutting down and reseeding the plots. No long-term changes in management with stand age were reported by the farmers in our study; however, it is likely that as plants matured, the frequency of management increased as harvesting became possible. We also found that like bacterial richness, the concentration of calcium and potassium decreased with plant age. Although these were not correlated to each other, both calcium and potassium were strong drivers of bacterial community composition. Importantly, both fungal and bacterial richness decreased over time. Thus, it is possible that the gradual decrease in richness is a result of the continued pressures of management on the soil microbiota.

Our fungal functional guild analyses revealed that endophytic fungi made up significant proportions of the fungal communities across all coffee farms regardless of management, previous land use or time since cultivation. Fungal genera categorized as endophytes are generally nonpathogenic endosymbionts, forming symbioses with host plants in which endophytes can provide protective benefits against pathogenic fungi, or simply coexist as commensals (Wilson 1995). Our findings are consistent with previous studies documenting abundant and diverse communities of endophytic fungi in coffee agroecosystems, but most studies examined fungal endophytes in the phyllosphere of *Coffea* plants (Vega *et al*. 2010; Oliveira *et al*. 2014; Saucedo-García *et al*. 2014). The relative abundances of endophytes across management categories in our study showed a trend of decreasing relative abundances of endophytes under conventional management compared to organic. Further, our results show a trend in the opposite direction for fungal plant pathogen communities, in which there was higher relative abundance of plant pathogens in conventionally managed soils compared to organic. This finding is likely due to a combination of biotic and abiotic drivers of fungal community assembly. First, interactions between populations of endophytes and pathogens may be driven by biotic interactions in which endophytes protect plants against antagonistic fungi known to cause disease; indeed, these interactions have been briefly examined in coffee systems (Vega *et al*. 2010). Another way in which management may drive shifts in the relative abundances of pathogenic fungal communities is through differences in abiotic conditions. The relationship we show here between relative abundances of fungi that cause disease in plants, and the soil chemical environment may be indirectly driven by management through fertilizer-induced shifts in the soil nutrient economy.

Our study highlights how multiple aspects of management alter the soil microbial communities in coffee agroecosystems. Previously, one study has shown how management affects soil nutrient availability and total microbial activity (Partelli *et al*. 2012), while another has found that microbial activity and biomass are better predictors of coffee productivity than soil chemistry (Silva Aragão *et al*. 2020). Similarly, we found that the soil microbiota covaried with several factors, including management, but most notably, previous land use and stand age. While we were not able to measure nutrient cycling rates, multiple studies have shown that biochemical processing rates, and decomposition in particular depend on the composition of the soil microbial community (Hartmann *et al*. 2015; Glassman *et al*. 2018; Ren *et al*. 2018). Thus, despite rarely being considered, it is likely that previous land use and stand age play an equally—if not more—important role in determining soil fertility for coffee production and the long-term sustainability of coffee agroecosystems.

## ACKNOWLEDGEMENTS

We would like to thank A. Gentile, C. Dean, and S. Tem for their help and thoughtful discussions, and M. Mark and C. Gillikin for help with permit acquisition. The authors declare no conflicts of interest.

