## Supplementary for "Soil microbial composition varies in response to coffee agroecosystem management"

SUPPLEMENTARY MATERIALS AND METHODS

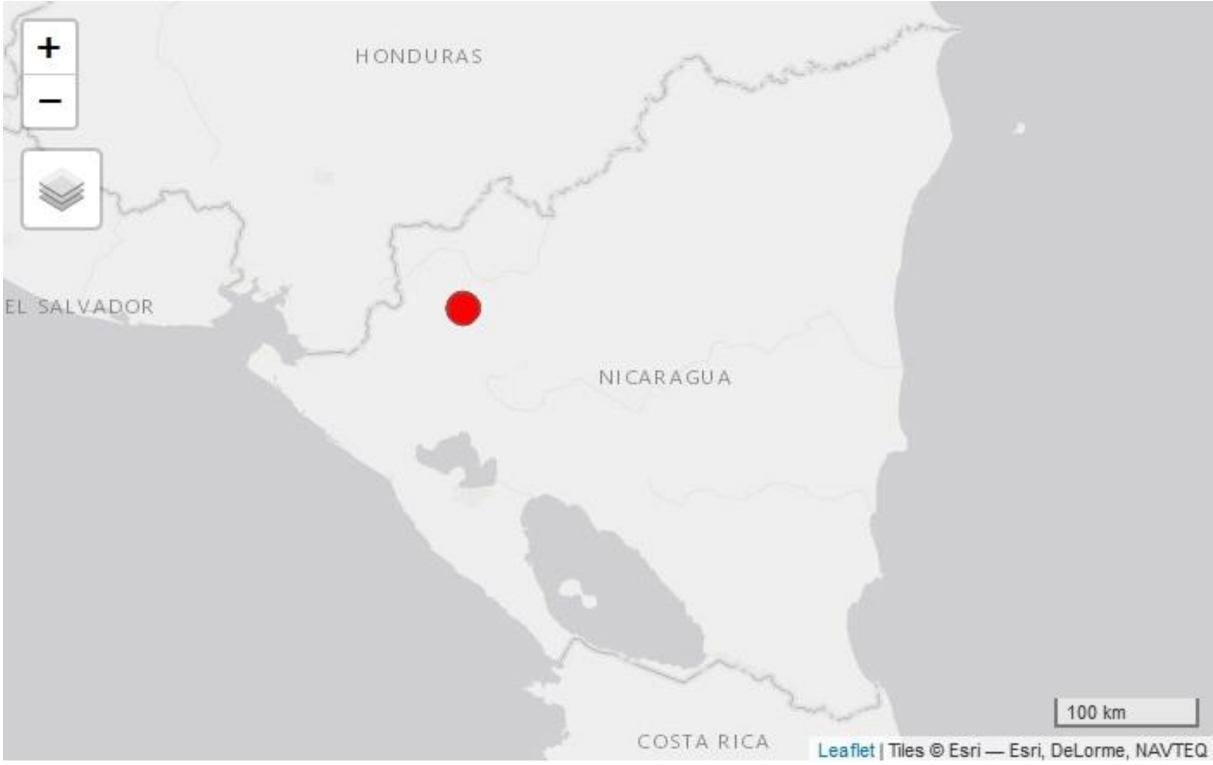


Figure S1. UCA Miraflor is located on the north-eastern corner of Nicaragua.

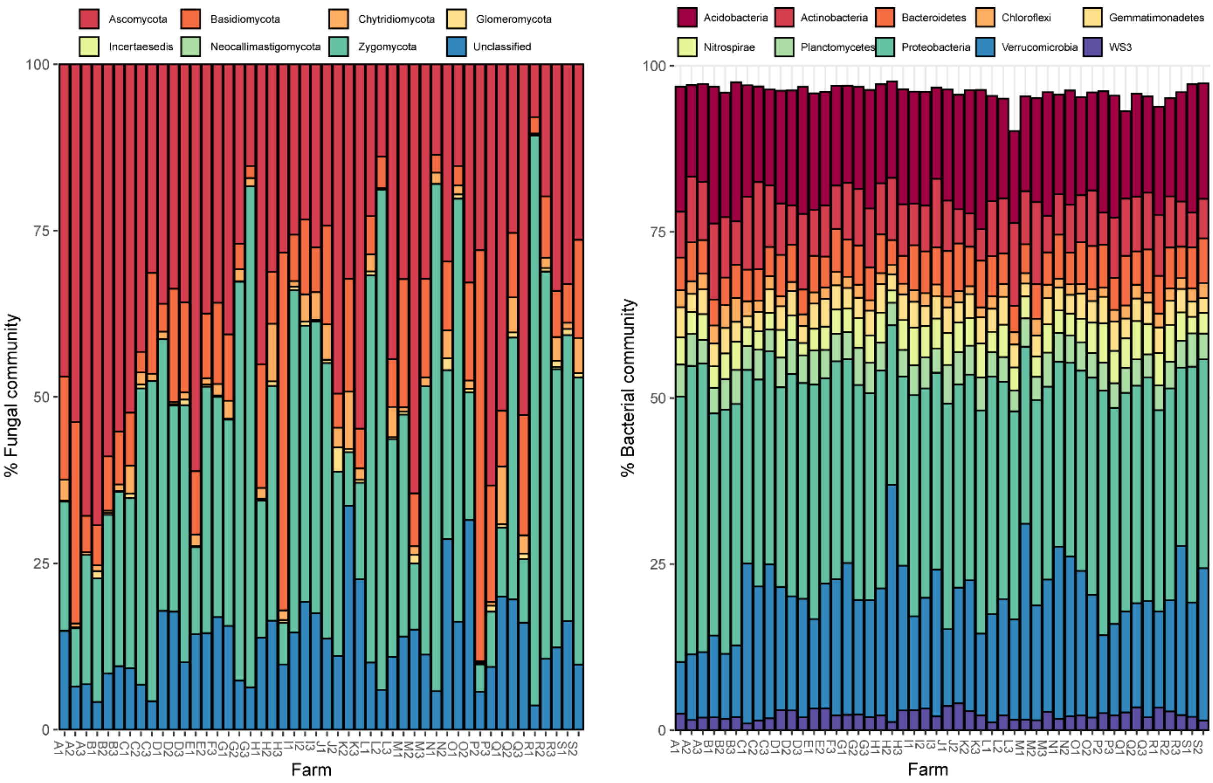


Figure S2. Relative abundance of the dominant fungal and bacterial phyla in each sample. For bacteria, the 10 most dominant phyla on average across all samples are displayed. All fungal phyla are represented.

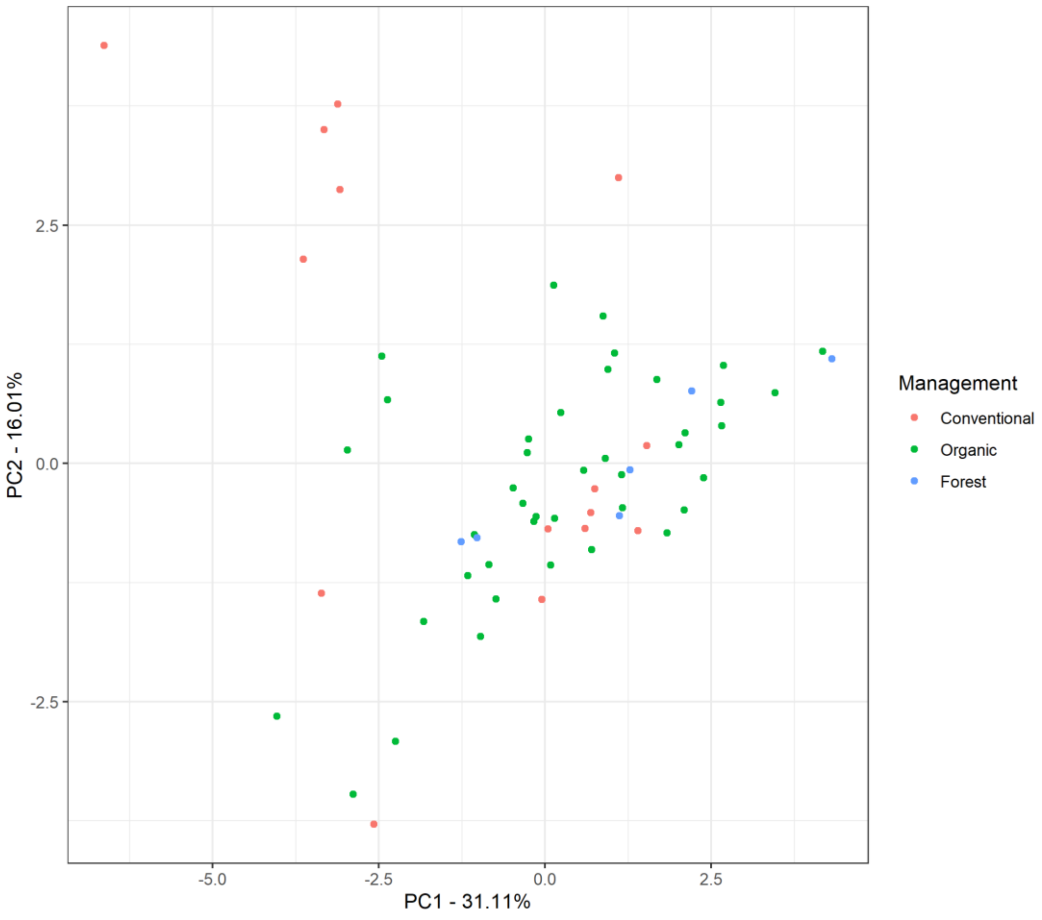


Figure S3. Biplot of principal component axes 1 and 2 for soil chemical variables Ca (ppm), K (ppm), Mg (ppm), P (ppm), Al (ppm), Ba (ppm), Fe (ppm), Mn (ppm), Na (ppm), Zn (ppm), B (ppm), Cu (ppm), total %C, total %N, and moisture content (%). Points represent individual samples, highlighted by management.

Table T1. Characteristics of each of the farms sampled in this study.

| **Farm code** | **Coffee age (years)** | **Previous land use** | **Management** |
|---|---|---|---|
| **A** | 4 | Primary | Conventional |
| **B** | 10 | Coffee | Conventional |
| **C** | 14 | Secondary | Organic |
| **D** | 45 | Coffee | Organic |
| **E** | 8 | Coffee | Organic |
| **F** | 10 | Secondary | Conventional |

| | | | |
|---|---|---|---|
| **G** | 12 | Secondary | Organic |
| **B** | 13 | Secondary | Organic |
| **I** | 11 | Primary | Organic |
| **J** | 10 | Primary | Organic |
| **K** | 2 | Primary | Organic |
| **L** | 13 | Secondary | Organic |
| **M** | 15 | Coffee | Conventional |
| **N** | 10 | Primary | Conventional |
| **O** | 40 | Coffee | Organic |
| **P** | 1 | Secondary | Organic |
| **Q** | 1 | Coffee | Organic |
| **R** | 4 | Coffee | Organic |
| **S** | 10 | Secondary | Organic |
